# TolC is required for a Mixed-Linkage β-Glucan (MLG) biosynthesis: Engineering bacteria for MLG overproduction

**DOI:** 10.64898/2026.04.30.721817

**Authors:** L. Ruiz-Sáez, P.J. Pacheco, J. Peinado, J. Lloret, S. Muñoz, J. Sanjuán, D. Pérez-Mendoza

## Abstract

Mixed-linkage β-glucans (MLGs) are emerging as promising biopolymers with significant biotechnological potential due to their unique structural and rheological properties. In rhizobia, MLG biosynthesis is controlled by the second messenger cyclic di-GMP (c-di-GMP) and mediated by the bicistronic operon *bgsBA*. However, the full composition of the biosynthetic machinery and strategies for enhanced production remain incompletely understood. In this study, we demonstrate that the outer membrane protein TolC is essential for MLG production in *Sinorhizobium meliloti*. Genetic disruption of *tolC* abolished MLG synthesis, while its complementation restored production. We propose that TolC forms a tripartite complex with BgsA and BgsB, enabling efficient polymer synthesis and export. Furthermore, co-overexpression of *tolC, bgsBA*, and a constitutively active diguanylate cyclase (*pleD\**) yielded a 10-fold increase of MLG over a control plasmid without *tolC*, reaching up to ∼10 g/L under bioreactor conditions. Additionally, this genetic module enabled de novo MLG production in otherwise non-producer rhizobial hosts (e.g. *Mesorhizobium japonicum*), allowing bacterial chassis exchanges and highlighting its portability and potential for synthetic biology applications. Overall, our findings identify TolC as a key component of the MLG biosynthetic machinery and provide a robust platform for the scalable production of this valuable biopolymer.

**Graphical Abstract:** 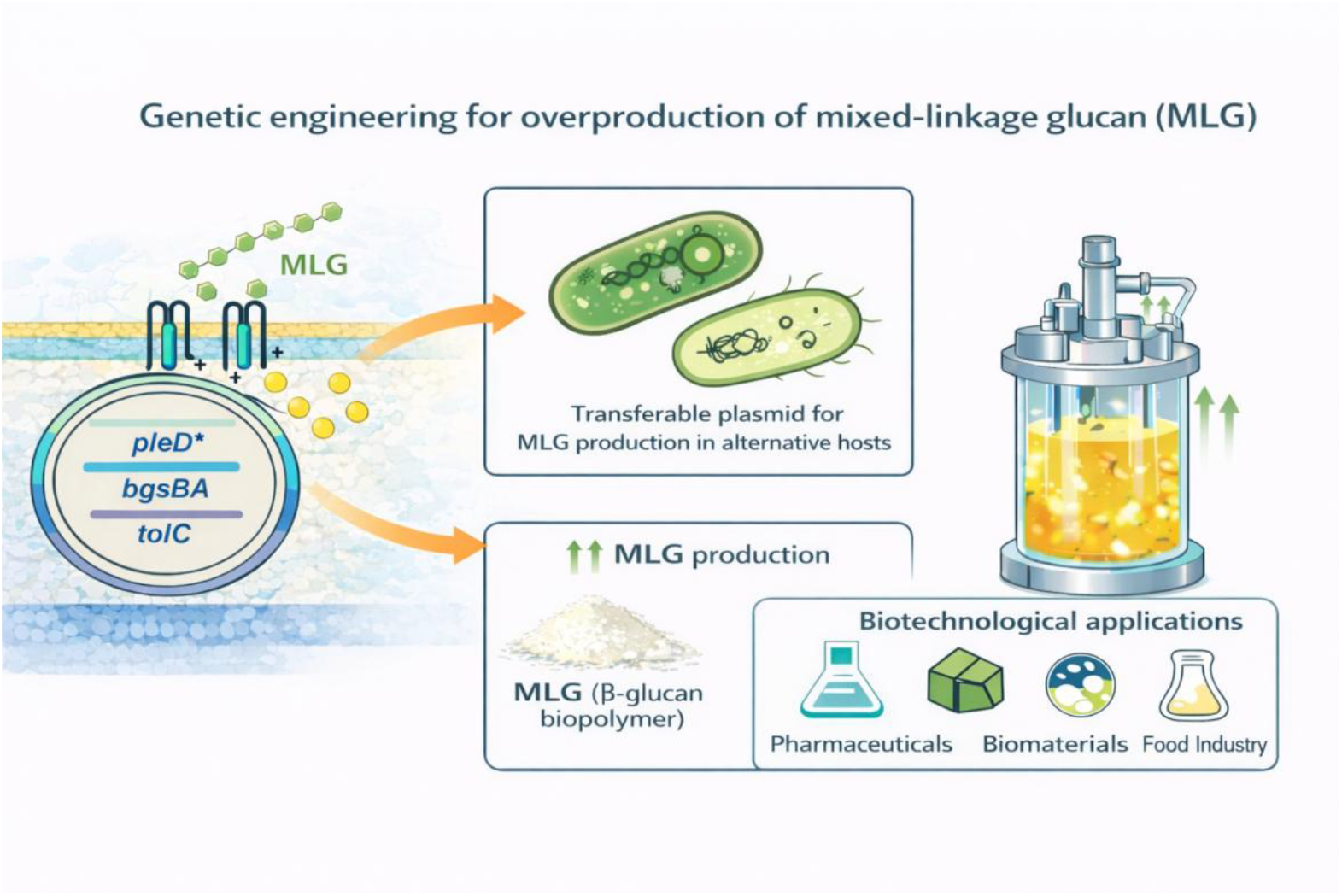

## Introduction

Bacterial exopolysaccharides (EPS) have increasingly captured scientific attention due to their significant commercial relevance in industries such as pharmaceuticals, cosmetics, biotechnology, and biomedicine (Freitas et al. 2011). The versatility of bacterial EPS lies in their unique material properties, including solubility, rheological features, viscoelasticity, crosslinking, gelation, water retention, extensibility, ionic strength, and exceptional stability. These characteristics highlight their industrial potential and position them as promising alternatives to chemical polymers due to their biodegradable and non-toxic nature (More et al. 2014; Moradali and Rehm 2020). In addition, the ease of working with bacteria has made bacterial EPS preferable over their plant and fungal counterparts (Kaur and Dey 2023).

EPS biosynthesis is a complex intracellular process where sugar residues are polymerized and modified before being secreted (Angelin and Kavitha 2020). Three main pathways mediate EPS synthesis: Wzx/Wzy-dependent, ABC transporter-dependent, and synthase-dependent pathways. The latter, most common in Gram-negative bacteria, employs a membrane-embedded glycosyl transferase (GT) that facilitates the simultaneous formation and translocation of the polymer across the inner membrane (Whitney and Howell 2013). EPS production and secretion are tightly regulated processes governed by complex signalling networks that integrate cellular and environmental cues. Among the key regulatory mechanisms, the second messenger bis-(3′–5′)-cyclic dimeric GMP (cyclic diguanylate, c-di-GMP) has emerged as a ubiquitous bacterial signalling molecule and a central activator of EPS biosynthesis, acting at both transcriptional and post-translational levels (Liang 2015). This second messenger plays a key role in signal transduction pathways that regulate and enable bacterial adaptation to different changing environments, switching between the motile planktonic state and the sedentary lifestyle associated with biofilms (D’Argenio and Miller 2004; Hengge 2009; Römling et al. 2013). First identified by Benziman and colleagues in 1987 as an activator of cellulose synthase (CS) (Ross et al. 1987), c-di-GMP is now recognized as a widespread allosteric regulator of synthase complexes that promote the synthesis and secretion of diverse EPS (Pérez-Mendoza and Sanjuán 2016). Key proteins involved in c-di-GMP metabolism are diguanylate cyclases (DGCs) and phosphodiesterases (PDEs). DGCs, featuring a GGDEF domain, synthesize c-di-GMP from two GTP molecules, while PDEs, containing EAL or HD-GYP domains, degrade c-di-GMP into 5′-pGpG or GMP. The cellular c-di-GMP level is tightly regulated by the opposing activities of these enzymes, often modulated by their sensory domains (Tal et al. 1998; Römling et al. 2005).

Rhizobia are soil α-proteobacteria that establish nitrogen-fixing symbioses with leguminous plants. These bacteria exhibit remarkable adaptability, switching between free-living and symbiotic lifestyles in response to environmental conditions. Bacterial signal transduction serves as a crucial mechanism for generating the necessary physiological, genetic, and cellular adaptive responses to ensure survival and function under different circumstances. c-di-GMP signalling is essential to coordinate these transitions and regulates motility, biofilm formation and EPS production (Krol et al. 2020). In *Sinorhizobium meliloti* 8530, c-di-GMP modulates the synthesis of several EPSs, including EPS-I (succinoglycan) and EPS-II (galactoglucan), both essential for root colonization, nodule invasion and host signalling. Additionally, c-di-GMP promotes the biosynthesis of an arabinose-containing polysaccharide (APS) and an adhesive polymer of unknown composition that bears similarities to the unipolar polysaccharide (UPP) observed in *Agrobacterium tumefaciens* (Schäper et al. 2016; Schäper et al. 2017; Krol et al. 2020). Moreover, in *S. meliloti* c-di-GMP promotes the synthesis and secretion of MLG, a linear mixed-linkage β-glucan. Unlike plant-derived MLGs, this bacterial MLG features a unique primary structure with perfect alternation of β-(1,3) and β-(1,4) linkages, making it a β-glucan with promising biotechnological applications (Pérez-Mendoza et al. 2015). In *Rhizobium etli* CFN42, c-di-GMP has also been shown to regulate the biosynthesis of MLG, highlighting the conservation of this regulatory mechanism within different rhizobial species. This MLG exhibits structural and rheological characteristics that position it as a novel and promising smart material with strong potential for large-scale production in pharmaceutical, medical, and food-related applications (Chang et al. 2021; Caseiro et al. 2022; Flores et al. 2026). Its biosynthesis requires an intact quorum-sensing system (*expR, sinI* and *sinR*), suggesting cross-talk between cell density signalling and c-di-GMP pathways (Pérez-Mendoza et al. 2015). In addition to the biotech relevance of this polymer, this conservation underscores the important relevance of MLG in the rhizosphere and symbiotic contexts, possibly contributing to surface architecture, biofilm formation, and plant–microbe interactions. Interestingly, in *R. etli* CFN42, c-di-GMP not only promotes the synthesis and secretion of MLG but also cellulose (Pérez-Mendoza et al. 2022).

Biosynthesis of MLG is mediated by the bicistronic operon *bgsBA* (Pérez-Mendoza et al. 2015). Bioinformatic analyses indicate that BgsA is an integral membrane protein with seven predicted transmembrane domains, a cytoplasmic GT domain responsible for glucan polymerization, and a C-terminal domain capable of binding c-di-GMP. Binding of c-di-GMP to this C-terminal region activates MLG synthesis, a mechanism reminiscent of the role of PilZ domain in regulating cellulose synthase in other bacteria (Pérez-Mendoza et al. 2017). *BgsB*, encodes a membrane fusion protein (MFP) containing an HlyD-like domain. Such proteins are commonly associated with T1SSs, acting as periplasmic adaptors that bridge the inner membrane transporter with Outer Membrane Factors (OMF) to facilitate substrate extrusion from the cell (Pimenta et al. 2005; Zgurskaya et al. 2009). Regarding OMFs, TolC stands out as trimeric outer membrane protein organized in two main domains: a β-barrel pore in the outer membrane and an α-helical domain projecting inward, where it interacts with periplasmic adaptors containing HlyD domains (Lee et al. 2012).

Similar to other β-glucans, like cellulose or curdlan, whose biosynthetic machineries involve several inner and outer membrane proteins, it would be expected that MLG biosynthesis require additional proteins besides BgsA and BgsB. The aims of this work were (i) to identify new, yet unknown components of the MLG biosynthesis machinery, and (ii) to engineer bacteria for increased MLG production. We demonstrate that the outer membrane porin TolC is required for MLG production, likely forming with BgsBA a tripartite complex capable of synthesizing and exporting MLG out of the cell. Moreover, its combined overexpression with *bgsBA* and *pleD*^*\**^ enables not only enhanced polymer yields but also *de novo* synthesis in alternative, naturally non-producer bacterial chassis.

## Experimental procedures

### 2.1 Bacteria and Culture Conditions

#### Bacterial strains and plasmids used in this study are listed in Table S1

*Sinorhizobium (Ensifer) meliloti* and *Rhizobium etli* were routinely grown in Tryptone-Yeast extract-CaCl_2_ (TY) (Beringer 1974) or in Minimal Medium (MM) (Robertsen et al. 1981) at 28-30ºC. When required, antibiotics and other compounds were added at the following final concentrations: Tetracycline (Tc), 10 µg/ml (5 µg/ml for *R*.*etli*); Kanamycin (Km), 50 µg/ml; Congo Red (CR), 50 µg/ml; Calcofluor White (CF, Sigma-Aldrich (o la marca correcta)), 200 µg/ml; IPTG, 1 mM. All the plasmids used in this study (Table S1) were introduced by conjugation using *E. coli* β2163 or S17.1 donor strains.

### 2.2 MLG production and quantification in flasks

MLG was quantified by measuring glucose equivalents released after lichenase digestion, following the methodology described by Marchante *et al*., (Marchante et al. 2024). Rhizobia were grown in 100 mL Erlenmeyer flasks containing 20 mL of liquid Minimal Medium supplemented with Tc at the appropriate concentration. Cultures were incubated at 28-30°C for 3 days with shaking (170 rpm), starting with an initial OD_600_ of 0.05, achieved by resuspending colonies from solid medium.

Cultures with flocs were transferred to 15 mL Falcon tubes and centrifuged at 3739 x g for 30 minutes at room temperature. The supernatant was discarded, and the resulting pellet was washed twice (20 min per wash) with 5 mL of 100 mM sodium phosphate buffer (pH 6.8). The washed flocs were subsequently resuspended in 1 mL of 100 mM sodium phosphate buffer and transferred to 2 mL Eppendorf tubes, for treatment with 6 units of lichenase enzyme (Megazyme, EC Number: 3.2.1.73). Enzymatic digestions were carried out at 40°C for 30 minutes with constant agitation. After digestion, the tubes were centrifuged at 9290 x g for 10 minutes, and the supernatant was collected for subsequent sugar quantification.

Soluble sugars were quantified using the method described by (Masuko et al. 2005). Briefly, 50 µL of the supernatant previously obtained were transferred to a multi-well plate, adding 150 µL of 97% (w/w) sulfuric acid and 30 µL of 5% (v/v) phenol, in that order. The plate was incubated at 90°C for 15 minutes and then placed in a microplate reader (BioTek®) to measure absorbance at 490 nm. Simultaneously, a calibration curve was constructed using different glucose dilutions to calculate glucose-equivalent concentrations in each sample.

### 2.3 Protein quantification by Bradford assay

Total protein concentration was determined according to Bradford (Bradford 1976) to normalize the results obtained in sugar quantification. Briefly, cell pellets obtained from 20 ml cultures were resuspended in 300 µl PBS buffer, pH 6.8-7.0. An aliquot of 20 µL from each suspension was diluted 1:1 with sterile distilled water and mixed with 160 µL of Bradford reagent (Bio-Rad). After 5-10 min of incubation at room temperature, absorbance was measured at 595 nm using a microplate reader. Protein content was calculated using a bovine serum albumin (BSA) standard curve (from 0 to 40 µg/mL), correcting for the dilution factor, and expressed as total mg per pellet.

### 2.4 Molecular cloning

The *tolC* gene from *S. meliloti* was amplified using primers TolC-F/TolC-R (Table S2) with an annealing temperature of 62°C and an extension time of 90 seconds. The resulting product (1457 bp) was cloned into the pCR2.1-TOPO® vector (Invitrogen) and confirmed by sequencing. The *tolC* fragment was then digested with *Bam*HI and ligated into the *Bam*HI-linearized pBBR9091 vector (Table S2), which harbors the *bgsBA* operon. Insert orientation was confirmed by *Mlu*NI digestion. Subsequently, the *bgsBA-tolC* fragment was excised from pBBR by *Xba*I digestion and ligated into the *Xba*I-linearized pJB*pleD*\* vector previously treated with 1 µL (5 U) Antarctic Phosphatase (New England Biolabs) according to the manufacturer’s instructions to prevent vector self-ligation. The correct orientation of the insert was confirmed by *Bam*HI digestion, obtaining the plasmid pJB*pleD*\*9091C.

To generate plasmid pJB*pleD*\**lacI*^q^, a 1610 bp fragment containing the *lacI*^*q*^ gene was excised from pQE-80L by *Mlu*NI digestion, purified and ligated with pJB*pleD*\*, which was previously linearized with *Bsa*AI and treated with Antarctic Phosphatase (NEB) as previously mentioned… The ligation mixture was transformed into chemically competent *E. coli* DH5α cells. Correct plasmid construction was verified by *Xba*I/*Eco*RI double digestion, yielding fragments of approximately 6200 bp, 2400 bp, and 1400 bp, consistent with the expected recombinant plasmid.

In-frame gene deletions were carried out in *S. meliloti 8530* and IBR505 following the protocol described by Schäfer *et al*. (Schäfer et al. 1994). Briefly, a synthetic DNA containing approximately 750 bp of each gene-flanking region was cloned into the suicide plasmid pK19*mobsacB*. After conjugation, neomycin-resistant colonies were selected as evidence of plasmid integration into the genome. These colonies were subjected to sucrose selection. The mucoid EPS II phenotype and the Calcofluor fluorescence EPS I phenotype were evaluated for colony selection. Successful gene deletion was confirmed by PCR analysis. The deletions targeted the following genes: *wgcA* (EPS II), *exoY* (EPS I) and *uxs1-uxe* (APS),and were obtained sequentially in that order as no antibiotic resistance marker was used.

### 2.6 Random mutagenesis strategy

To generate random transposon mutants in *S. meliloti* 8530 and IBR607 strains, the transposon Tn*5* (which confers Km/Sm resistance and is carried by plasmid pSUP2021, Table S1) or the transposon TnSAM (which confers Km resistance and is carried by pSAM_Rl, Table S1) were mobilized from *E. coli* donor strains S17.1 or β2163, respectively, by conjugation following the filter-mating procedure with a donor:recipient ratio of 1:1 (Demarre et al. 2005). Transposants were cultured on TY broth supplemented with neomycin or kanamycin. For *S. meliloti* 8530 mutagenesis, plasmid pJB*pleD*\**lacI*^q^ (encoding an exogenous diguanylate cyclase from *Caulobacter crescentus*, Table S1) was transferred from β2163 to the pool of transposon mutants. Transconjugants were selected on TY or MM plates supplemented with CR, IPTG and Tc. In parallel, *S. meliloti* 8530 strain harbouring pJB*pleD*\**lacI*^q^ or pJB3Tc19 plasmid was plated onto CR-containing medium also supplemented with Tc and IPTG as positive or negative control, respectively. The CR-negative^−^ colonies obtained from both Tn5 and TnSAM mutagenesis were isolated and their transposon insertion sites were determined by random PCR according to Saavedra *et al*. (Saavedra et al. 2017) or by whole-genome sequencing performed by Plasmidsaurus using Oxford Nanopore Technology. Different primer pairs were designed and used depending on the transposon (Tn*5* or TnSAM-harbouring colonies) and the inverted repeat region to amplify (IRR or IRL) (Table S2). The amplified PCR products were purified from agarose gels and sequenced using primers Tn5F2 for Tn5 and IRL3 or SAMIRRSEC for TnSAM (Table S2). DNA sequence homology searches were performed with the BLAST program from NCBI to determine the precise positions of the transposon insertions.

### 2.7 Bioreactor culture conditions for MLG production

*S. meliloti* cultures were scaled up progressively. Initially, cells were inoculated into 20 mL of TY broth supplemented with Tc and incubated for 24 h at 28 °C (OD_600_ = 1.0). This pre-culture was transferred to 50 mL of the same medium and incubated for another 24 h under identical conditions. Subsequently, this culture was used to inoculate a 2.5 L bioreactor at a 2 % (v/v) concentration. The laboratory-scale bioreactor contained minimal medium (MM; (Robertsen et al. 1981) with the following composition (per litre): 10 g mannitol, 1.1 g sodium glutamate, 0.3 g KH_2_PO_4_, 0.3 g K_2_HPO_4_, 0.15 g MgSO_4_·7H_2_O, 0.05 g CaCl_2_·2H_2_O, 0.006 g FeCl_3_, and 0.05 g NaCl. Additionally, 1 mL of a 1000× vitamin solution (0.2 g/L biotin, 0.1 g/L thiamine hydrochloride, and 0.1 g/L sodium pantothenate) was added. The initial pH was adjusted to 6.8– 7.0. The total fermentation time was 96 h, with a fed-batch addition of 0.5 L of the same medium after 72 h to provide additional nutrients.

Dissolved oxygen (DO) levels were maintained above 95% saturation using a cascade control system with two priority parameters: agitation speed (800–1000 rpm) and aeration rate (0.2–0.4 slpm). Throughout the process, the pH was maintained between 7.0 and 8.0 by the automatic addition of 2 M H_2_SO_4_ or 2 M NaOH, while the temperature was kept at 28 °C. To prevent foaming, an antifoam agent was added at a final concentration of 0.01% (v/v).

### 2.8 Extraction and purification of MLG

After 72 h of fermentation, the culture was distributed into 0.5 L bottles and centrifuged at 14,000 × g for 30 min. The supernatant was filtered to recover any released flocs. Pellets were then resuspended in Milli-Q water and centrifuged again under the same conditions to ensure the complete removal of residual medium. This step was repeated until a single MLG-containing pellet was obtained. The resulting pellet (MLG plus cells) was boiled for 10 min at 100 °C and washed with Milli-Q water. This washing step was repeated three times. The purified MLG was then transferred into a dialysis membrane (6-8 kDa molecular weight cut-off) and dialyzed against 3 L of Milli-Q water for 24-48 h, with the water being replaced after the first 24 h. Finally, the dialyzed MLG was frozen at −80 °C and lyophilized for 24 h. The resulting dry MLG was weighed to determine the yield.

## Results

### Effect of other c-di-GMP–regulated EPS on MLG production in rhizobia

Intracellular c-di-GMP levels regulate the synthesis of multiple EPS in *S. meliloti* 8530 (Krol et al. 2020). With the aim of maximizing MLG production, we constructed a triple mutant defective in the synthesis of three c-di-GMP–regulated EPS in this bacterium: EPS II, EPS I, and APS. To this end, in-frame deletions were generated in the following genes: *wgcA* (EPS II), *exoY* (EPS I), and *uxs1–uxe* (APS). These mutations were introduced sequentially and confirmed both genetically by PCR and microbiologically by the absence of the phenotypes associated with each EPS. Once this triple mutant (IBR606) had been constructed, the MLG yield of this mutant was compared with that of the wild type (Fig. 1A). In line with previous reports, at physiological c-di-GMP levels MLG production in the wild type remained below the detection limit, and no relevant increase was observed in the IBR606 background (Fig. 1A). As expected, the presence of plasmid pJB*pleD*\* boosted MLG production in IBR606, which was slightly higher than in the wild type, although this difference was not statistically significant (Fig. 1A). These results indicate that the ability of *S. meliloti* to synthesize other c-di-GMP-regulated EPS, such as EPS I, EPS II and APS, does not have a major impact on MLG yield.

**Figure 1.**
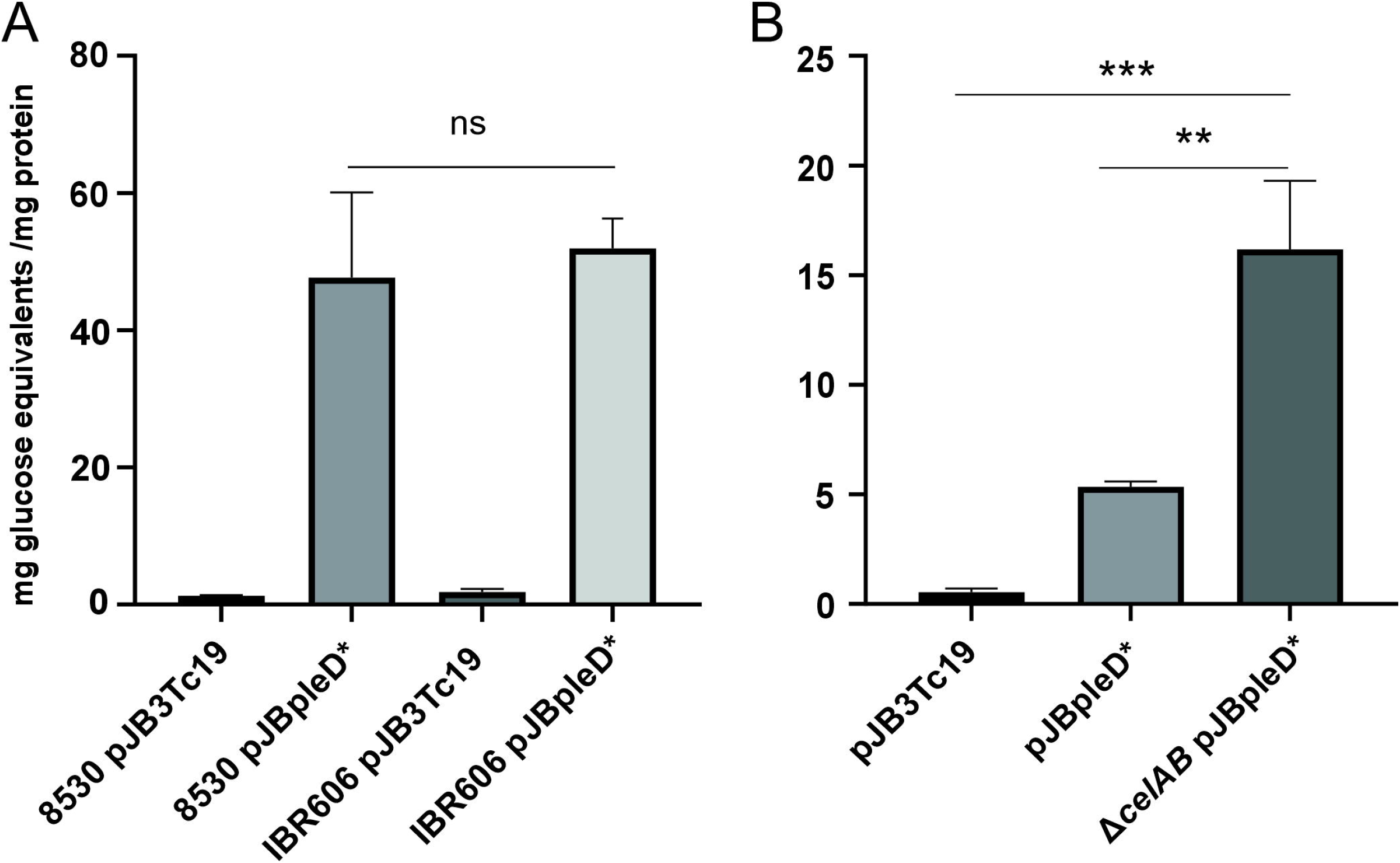
Effect of other c-di-GMP–regulated EPS on MLG production. Quantification of MLG in *Sinorhizobium meliloti* (A) and *Rhizobium etli* (B). Data were normalized to total protein content and represent the mean ± SD of independent experiments (n = 8 for *S. meliloti*; n = 4 for *R. etli*). In panel A, ns = no statistically significant differences according to a two-tailed Welch’s unpaired t-test. In panel B, statistical significance: according to a one-way ANOVA followed by a Tukey’s multiple comparisons test: \*\**p* < 0.01; \*\*\**p* < 0.001.

A similar analysis was performed in *R. etli* CFN42, which is another MLG-producer where c-di-GMP activates the production of both MLG and bacterial cellulose, two β-glucans that utilize UDP-glucose as a substrate (Pérez-Mendoza et al. 2022). MLG production was compared between the wild type strain and a cellulose-deficient mutant (Δ*celAB*). In contrast to what was observed in *S. meliloti*, under elevated c-di-GMP conditions (pJBpleD*) the *ΔcelAB* mutant exhibited a significant ∼3-fold increase in MLG production relative to the wild type (Fig. 1B). These results indicate that blocking cellulose production is an effective strategy to increase MLG yield in the *R. etli* genetic background. However, as shown in Figure 1, the *S. meliloti* 8530 background remains a superior bacterial chassis for MLG production compared to *R. etli ΔcelAB* (∼50 mg glucose equivalents/mg protein in the former vs ∼15 mg glucose equivalents/mg protein in the latter).

### Identification of Genes Potentially Involved in MLG Synthesis and Secretion in *Sinorhizobium meliloti*

To further optimize MLG production in *S. meliloti*, we investigated whether additional, yet unidentified genes contribute to MLG synthesis and secretion. Other β-glucans, such as cellulose and curdlan, rely on auxiliary components for full functionality (Stasinopoulos et al. 1999; Whitney and Howell 2013). To this end, we employed a random transposon mutagenesis approach followed by screening on solid medium supplemented with CR, which binds to α-D-glucopyranosyl units, neutral or basic polysaccharides, and certain proteins, and has been shown to interact positively with MLG (Pérez-Mendoza et al. 2015). The goal was then to distinguish between MLG-producing (red-stained) and -non-producing (white) colonies on CR plates after the mutagenesis. To maximize screening success, we applied this approach across different variants: (i) two different *S. meliloti* 8530 genetic backgrounds (wild-type or a EPS II, EPS I, and APS triple mutant), (ii) two different transposons (Tn*5* or TnSAM), and (iii) two different sources of c-di-GMP: via heterologous overexpression of *pleD\** or activation of the cognate DGC BgrR (Baena et al. 2019).

The first approach entailed a random transposon mutagenesis of the *S. meliloti* 8530 wild-type strain by introducing the Tn*5* transposon using the donor strain *E. coli* S17.1 (pSUP2021) (Simon et al. 1983) or the TnSAM transposon using β2163 pSAM_Rl (Perry and Yost 2014). Approximately, 8.5 × 10^4^ Tn*5* and 1.6 × 10^5^ TnSAM transposon insertion mutants were obtained, respectively. Then, *pleD\** was transferred to each pool of transposants by conjugation from the β2163 pJBpleD*lacI^q^ donor strain. Transconjugants were selected on TY or MM plates supplemented with CR, Tc and IPTG. In parallel, the non-mutagenized *S. meliloti* 8530 wild-type strain harbouring either pJBpleD*lacI^q^ or pJB3Tc19 plasmid was plated onto the same selective media as positive and negative controls, respectively. All non-mutagenized colonies appeared red (pJBpleD*lacI^q^) or white (pJB3Tc19), respectively. In contrast, some transposant colonies (<1%) of *S. meliloti* 8530 mutagenized with Tn*5* or TnSAM exhibited a CR-negative phenotype (white). Eight of these colonies, 3 with Tn*5* and 5 with TnSAM, were subjected to further investigation to identify the genomic sites of transposon insertion (Table 1). All transposon insertions were found interrupting genes previously described to be involved in MLG production. Two insertions were mapped to the *expR* gene (*Smc03896*), a transcriptional regulator required for the expression of the MLG biosynthetic operon *bgsBA* (Pérez-Mendoza et al. 2015), while 4 and 2 insertions were located within the *bgsB* and *bgsA* synthesis genes, respectively (Table 1).

**Table 1.** Mapping Transposon insertions. Results obtained from sequencing after random PCR on genomic DNA from transposon Tn5- or SAM-harbouring CR^-^ colonies.

| Genetic background | c-di-GMP source | Transposon used | Targeted gene | Number of insertions and their genome position* |
| --- | --- | --- | --- | --- |
| Sm 8530 | PleD* | Tn5 | *bgsB* (Smb20390) | 2 (403235 and 403073 of the pSymB) |
|  |  |  | *expR* (Smc03896) | 1 (3566930 of the chromosome) |
|  |  | TnSAM | *bgsB* (Smb20390) | 2 (402524 and 403054 of the pSymB) |
|  |  |  | *bgsA* (Smb20391) | 2 (404024 and 404772 of the pSymB) |
|  |  |  | *expR* (Smc03896) | 1 (3566858 of the chromosome) |
| Sm IBR607 | BgrR | TnSAM | *bgsB* (Smb20390) | 3 (943, 1139, ND) |
|  |  |  | *bgsA* (Smb20391) | 4 (1165, 1491, 1897, ND) |
|  |  |  | *expR* (Smc03896) | 4 (-1, 9, 368, 655) |

\*ND: exact position not determined

After failing to identify new genes involved in MLG production using the initial mutagenesis strategy, we decided to repeat the approach in an alternative *S. meliloti* 8530 genetic background. On one hand, we aimed to minimize potential interference from the synthesis of other EPS (using a triple mutant lacking EPS II, EPS I, and APS), and on the other, to modify c-di-GMP-dependent activation by using a native DGC (BgrR) specifically dedicated to MLG production. For this purpose, we used mutant IBR505, which lacks the *bgrV* gene, which encodes a repressor of the DGC BgrR, and therefore this mutant displays derepressed MLG synthesis (Baena et al. 2019). Sequential allelic exchange was then used to introduce deletions in *wgcA* (EPS II), *exoY* (EPS I), and *uxs1–uxe* (APS) into this genetic background, yielding the quadruple mutant IBR607 (defective in EPS II, EPS I, APS, and *bgrV*). IBR607 was subjected to mutagenesis with the mariner transposon pSAM_Rl, generating approximately 6,000 transposants. Following screening, 11 colonies showing a CR-negative phenotype were selected, and the respective transposon insertion sites were determined. As summarized in Table 1, the transposon was found to disrupt again any of the genes already known to be required for MLG production: *expR, bgsB*, and *bgsA*.

In summary, both approaches repeatedly recovered mutations in previously known genes, suggesting two possibilities: (i) no additional genes participate in MLG synthesis and secretion, or (ii) such genes do exist but are essential for *S. meliloti* under our experimental conditions.

### Role of TolC in MLG production

Other β-glucans, such as cellulose and curdlan, rely at a minimum on OMFs to mediate EPS extrusion into the extracellular milieu. In this context, TolC has been described as a highly versatile porin involved in several transport systems responsible for exporting diverse molecules and environmental stressors, including a broad range of antimicrobial agents (Wright et al. 2025). Notably, in *S. meliloti*, TolC is essential for the proper synthesis and secretion of the two symbiotically relevant EPS produced by this strain, namely succinoglycan (EPS-I) and galactoglucan (EPS-II). Furthermore, an insertional mutation in *tolC* in this strain severely compromises resistance to antimicrobials and increases susceptibility to osmotic and oxidative stresses (Cosme et al. 2008). Taken together, these observations led us to ask: (i) whether, as described for other *S. meliloti* EPS, TolC might also play a relevant role in MLG synthesis and/or secretion, and (ii) whether the failure to recover *tolC* in our random mutagenesis screens could be explained by the lack of viability of *tolC* mutants on CR–supplemented medium.

To address this, we first examined the growth capacity of *tolC* mutants of *S. meliloti* 1021 and 2011 strains on our screening medium supplemented with CR. Unlike their parental wild-type strains, these mutants showed no appreciable growth on MM or TY plates containing 50 µg/ml CR (data not shown). These results suggest that TolC is indeed essential under our screening conditions and prompted us to assess MLG production in these mutants. To this end, we introduced plasmid pJBpleD*9091, which carries *pleD\** and *bgsBA* genes under the control of a constitutive *lac* promoter (Pérez-Mendoza et al. 2015), into the 1021 or 2011 *tolC* mutant backgrounds. Unlike *S. meliloti* 8530 (*expR*^+^), strains 1021 and 2011 lack the transcriptional regulator ExpR and thus require the expression of *bgsBA* from a heterologous promoter for MLG production (Pérez-Mendoza et al. 2015). As shown in Fig. 2, plasmid pJBpleD*9091 allowed strains 1021 and 2011 to produce substantial amounts of MLG. In contrast, the *tolC* mutation in *S. meliloti* 1021 (Fig. 2A) and 2011 (Fig. 2B) genetic backgrounds resulted in a complete loss of MLG production. To corroborate the requirement of TolC for MLG production, we constructed a new plasmid based on pJBpleD*9091 (*bgsBA* and *pleD\**), in which the *tolC* gene was additionally introduced into the same transcriptional unit under the control of the *lac* promoter, yielding pJBpleD*9091C (*bgsBA, pleD*\* and *tolC*). Complementation of both *tolC* mutants with pJBpleD*9091C plasmid restored MLG production. These results demonstrate that TolC is required for efficient MLG production by *S. meliloti*.

**Figure 2.**
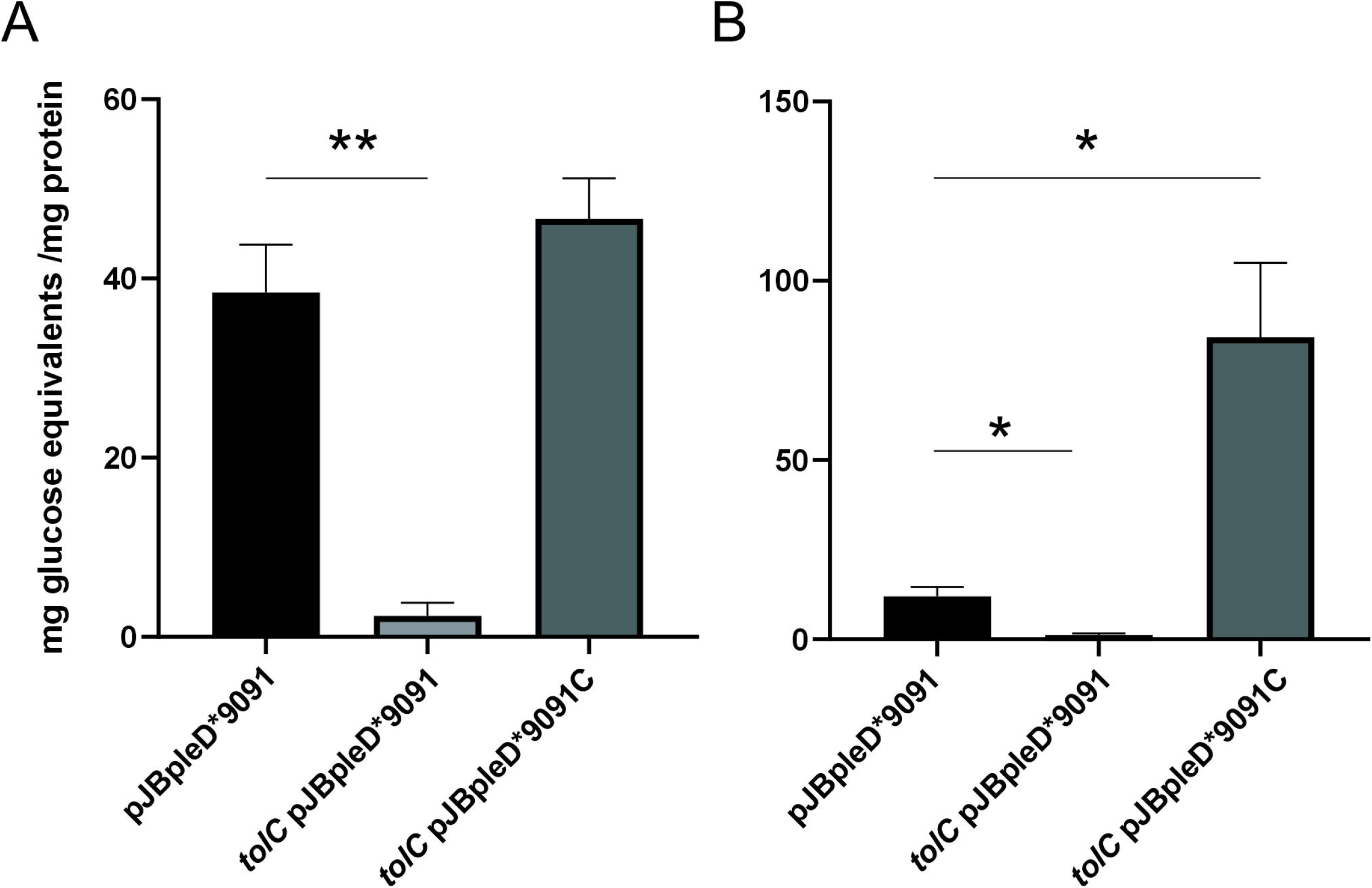
TolC is required for MLG production. Quantification of MLG production in *S. meliloti* strains 1021 (A) and 2011 (B) and their *tolC* mutant derivatives. MLG levels were measured in the wild-type strains and their corresponding *tolC* mutant carrying the pJBpleD*9091 plasmid (*bgsBA* and *pleD\**), as well as in the *tolC* mutant complemented *in trans* with *tolC* (pJBpleD*9091C). Data were normalized to total protein content and represent the mean ± SD from five independent experiments. Statistical significance between strains was determined using a two-tailed Welch’s unpaired *t*-test: \**p* < 0.05, \*\**p* < 0.01.

### TolC Overexpression enhances MLG production

The above complementation experiments with pJBpleD*9091C plasmid not only restored MLG production in the *tolC* mutants but also resulted in substantial increases over the wild-type levels, particularly in the 2011 *S. meliloti* background (Fig. 2B). From a biotechnological perspective, this finding is highly relevant for maximizing MLG yield. To confirm the enhancing effect of *tolC* overexpression on MLG production, we measured MLG levels in the *S. meliloti* 8530 Δ*bgsBA* background carrying the pJB3Tc19, the pJBpleD*9091 or the pJBpleD*9091C plasmids (Fig. 3A). As shown in Figure 3B, multicopy *tolC* expression yielded a significant ∼25 % increase in MLG production—the highest levels measured in our laboratory to date, exceeding 250 mg glucose equivalents per mg of protein.

**Figure 3.**
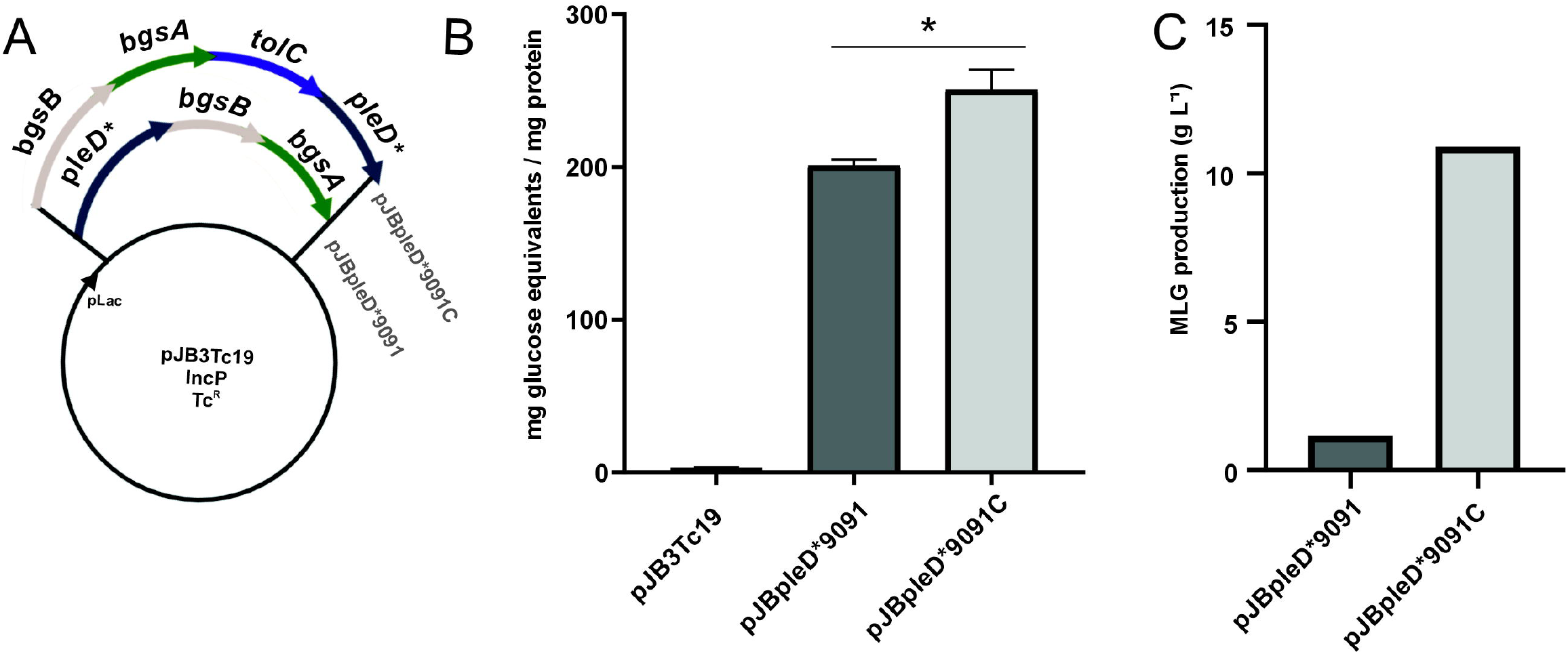
TolC overexpression increases MLG yields in *S. meliloti*. (A) Plasmid structure of pJBpleD*9091 and its derivative pJBpleD*9091C overexpressing *tolC*, constructed in this work. (B) MLG production was quantified in the *S. meliloti* 8530 Δ*bgsBA* background harbouring pJB3Tc19, pJBpleD*9091, or pJBpleD*9091C plasmids. Data were normalized to total protein content and represent the mean ± SD from five independent experiments (n = 5). Statistical significance between pJBpleD*9091 and pJBpleD*9091C was determined using a two-tailed Welch’s unpaired *t*-test (\**p* < 0.05). (C) MLG yields obtained under biotechnological fermentation conditions (2.5-L laboratory bioreactor) with the Δ*bgsBA* background with native TolC levels (pJBpleD*9091) or overexpressing *tolC* (pJBpleD*9091C).

Having demonstrated the enhancing effect of *tolC* overexpression at flask scale, we evaluated its impact on MLG production at a larger, more controlled scale using a 2.5-L laboratory bioreactor. Throughout the process, dissolved oxygen was maintained above 95% saturation, pH was held stable at 7–8, and temperature was controlled at 28 °C, with agitation and aeration automatically adjusted to sustain these parameters. After a 96-h fermentation, MLG was isolated and purified following the methodology described previously (Pérez-Mendoza et al. 2015). Under these conditions, we compared MLG production in the *S. meliloti* 8530 Δ*bgsBA* background hosting pJBpleD*9091C (containing *tolC*) or the control plasmid pJBpleD*9091. Consistent with the indirect quantification of glucose equivalents (Fig. 3B), *tolC* overexpression resulted in nearly a 10-fold increase over the control, achieving ∼10 g/L of MLG (Fig. 3C). In summary, identifying the essential role of TolC and leveraging its overexpression has led to outstanding yields, paving the way for the future biotechnological exploitation of this promising biopolymer.

### Exploring MLG Production Across Distinct Genetic Background

The excellent MLG production levels obtained with the plasmid carrying the biosynthetic genes *bgsBA*, the c-di-GMP source (*pleD\**), and the *S. meliloti* porin *tolC* within a single transcriptional unit (pJBpleD*9091C; Fig. 3A), raised the question of whether this plasmid could be used to engineer alternative bacterial chassis to produce MLG. To address this, pJBpleD*9091C was introduced into two distinct backgrounds: (i) a *R. etli* derivative unable to produce cellulose and MLG (Δ*celAB* Δ*bgsA*, Pérez-Mendoza et al. 2022), and (ii) *Mesorhizobium japonicum* MAFF303099, a rhizobium that naturally lacks the *bgsBA* biosynthetic operon.

First, the resulting transconjugants were grown on different media supplemented with CF and compared to their corresponding negative controls carrying the empty vector pJB3Tc19 (Fig. 4A). CF binds to various pure and mixed-linkage β-glucans, including MLG and cellulose, and exhibits a characteristic blue-green fluorescence under long-wavelength UV light (Pérez-Mendoza et al. 2014). As shown in Figure 4A, the presence of pJBpleD*9091C induced clear fluorescence under UV illumination in both genetic backgrounds. Subsequent quantification in liquid MM medium confirmed that the Δ*celAB ΔbgsA* double mutant of *R. etli* and *M. japonicum* MAFF303099 are both capable of producing MLG in presence of pJBpleD*9091C, in contrast to their respective controls with the empty vector (Fig. 4B). From a biotechnological standpoint, these results open the possibility of exploring additional bacterial chassis for MLG overproduction, not only to maximize yield but also to exploit metabolic flexibility and optimize production costs.

**Figure 4.**
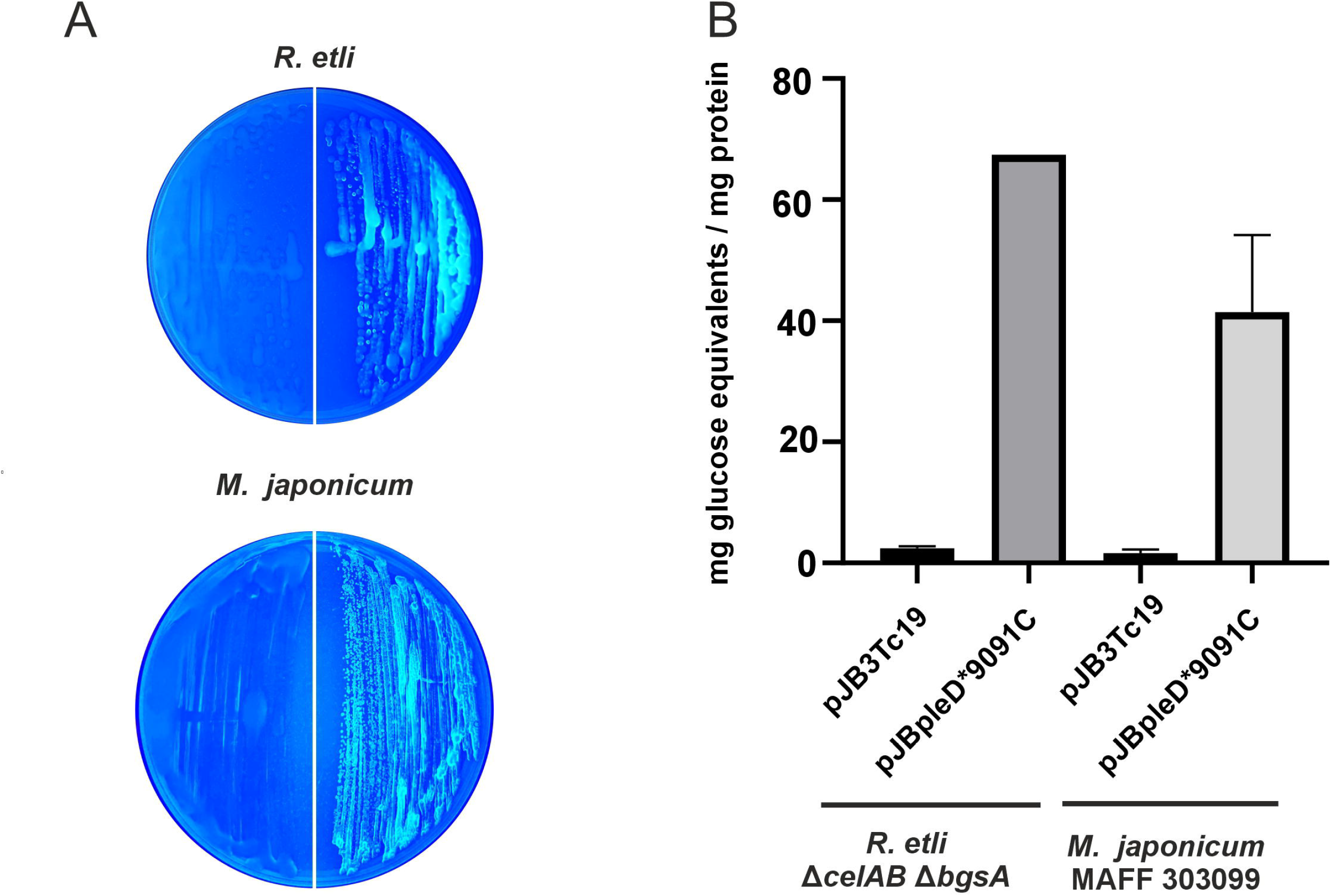
MLG production across distinct genetic backgrounds. MLG production in *R. etli* Δ*celAB* Δ*bgsA* and *M. japonicum* MAFF 303099 strains carrying pJBpleD*9091C or the empty vector pJB3Tc19. (A) Calcofluor phenotype of *R. etli* Δ*celAB* Δ*bgsA* and *M. japonicum* MAFF 303099 strains hosting pJB3Tc19 (left side of the plates) or pJBpleD*9091C (right side of the plates). Strains were grown on minimal medium supplemented with Calcofluor (200 µg mL^−1^) and photographed under UV illumination after 72 h. (B) Quantification of MLG in liquid cultures of both bacterial chassis containing pJB3Tc19 or pJBpleD*9091C. Data were normalized to total protein content and represent the mean ± SD from 3 independent experiments.

## Discussion

The second messenger c-di-GMP has been proposed in recent years as an important regulatory molecule controlling the production of bacterial EPS. This regulation can reach high levels of complexity in some bacteria, such as *S. meliloti*, where c-di-GMP can influence the biosynthesis of at least five different EPS (Krol et al. 2020). How this second messenger can regulate the biosynthesis of multiple EPS within the same bacterium remains a difficult question to answer. This question lies at the heart of the ongoing discussion about how c-di-GMP can simultaneously control a wide variety of processes in a given bacterium, e.g., motility, biofilm formation, cell cycle progression, etc… Recent studies establish that such high specificity and flexibility arise from the combination of local and global modes of c-di-GMP signalling within complex regulatory networks (Junkermeier and Hengge 2023). On EPS regulation, it could be explained by signalling systems that activate specific DGCs, which synthesize c-di-GMP in proximity to the biosynthetic complex involved in a given polysaccharide, thereby conferring spatial specificity to the system. From a biotechnological perspective, then, the situation could be completely different when a heterologous DGC (e.g. *pleD*\*) is artificially overexpressed with the aim of broadly raising the intracellular levels of c-di-GMP for increasing the production of a specific biopolymer of interest and no others (Pérez-Mendoza et al. 2026). With this in mind, and with the biotechnological goal of maximizing MLG production, we decided to investigate the potentially beneficial impact of funnelling metabolic flux toward MLG by blocking the synthesis of other c-di-GMP–regulated EPS in *S. meliloti*. However, the results in *S. meliloti* 8530 suggest that MLG biosynthesis is largely independent of other c-di-GMP regulated EPS pathways, despite the high metabolic cost of producing multiple polysaccharides simultaneously. The absence of a significant increase in MLG production in the triple mutant IBR606 (defective in EPS II, EPS I and APS) suggests that the biosynthetic flux toward MLG is sufficiently robust to accommodate normal resource allocation even in the presence of other active polysaccharide pathways. This finding aligns with previous reports indicating that MLG synthesis is regulated primarily by c-di-GMP rather than by competition for precursors among co-produced EPS (Pérez-Mendoza et al. 2014; Krol et al. 2020).

In stark contrast, in *R. etli* CFN42, MLG production appears to compete with bacterial cellulose biosynthesis. This effect can be attributed to the structural and metabolic similarities between cellulose and MLG, both β-glucans assembled by homologous glycosyltransferases from the same precursor (UDP-glucose) and both allosterically activated by c-di-GMP (Pérez-Mendoza et al. 2017; Pérez-Mendoza et al. 2022). Indeed, *R. etli* CFN42 appears to prioritize cellulose synthesis when c-di-GMP levels are high, likely due to the critical role of cellulose in biofilm formation and host plant colonization, which identified cellulose as the primary β-glucan contributing to these processes (Pérez-Mendoza et al. 2022). Only when cellulose production is abolished (*ΔcelAB* mutant) does MLG become overproduced in this genetic background, highlighting a hierarchy in β-glucan biosynthesis and suggesting that cellulose exerts a dominant pull on the shared metabolic precursors and regulatory signals. In any case, considering the MLG production levels achieved in both species, the genetic chassis of *S. meliloti* 8530 remains a superior platform for the hyperproduction of this c-di-GMP–dependent biopolymer. With the aim of further enhancing MLG production in this strain, we set out to identify new genetic factors that might be involved in MLG biosynthesis. The mutagenesis approaches used to identify additional genes involved in MLG production consistently yielded hits in *bgsA, bgsB*, and *expR*, supporting the idea that these genes constitute the core machinery for MLG synthesis and regulation. The absence of other candidates suggests either that the minimal MLG system relies primarily on these genes, or that essential components could not be recovered due to lethality or pleiotropic effects on the selective media used in this screening.

Among such essential components, TolC emerged as a compelling candidate. TolC is a well-characterized outer membrane protein and plays a critical role in T1SSs and multidrug efflux pumps, which are essential for the export of diverse metabolites and macromolecules (Koronakis et al. 1997; Nikaido 2011). This functional versatility further supports its potential involvement in the MLG biosynthetic machinery. Indeed, TolC homologs are generally required for the secretion of various EPS in Gram-negative bacteria, including *S. meliloti* (Cosme et al. 2008; Bina et al. 2023). This observation is particularly relevant given that different studies highlight the involvement of TolC-like proteins in c-di-GMP–regulated biological processes, such as biofilm formation, through the secretion of different adhesins via T1SSs (Monds et al. 2007; Pérez-Mendoza et al. 2011). In this context, our results show that *tolC* disruption in *S. meliloti* 1021 and 2011 completely abolishes detectable MLG levels, and that its production was restored when *tolC* was provided *in trans*. This intriguing result raises the question of whether the requirement for an intact *tolC* copy stems from the direct involvement of this porin in extruding the β-glucan chain through the outer membrane to the cell exterior forming a tripartite complex together with BgsB (HlyD) and BgsA (GT). Similar tripartite secretion systems have been reported for cyanobacterial extracellular cellulose production (Maeda et al. 2018; An et al. 2025). Furthermore, regarding other EPS, it seems reasonable to assume that MLG secretion requires an additional outer membrane component that interacts with BgsB to form a continuous channel through which MLG can be extruded outside the cell. Similar outer membrane porins have been described for other EPS, such as BcsC for cellulose, PgaA for poly-β-1,6-N-acetyl-D-glucosamine (pNAG), or AlgE for alginate (Tan et al. 2014; Wang et al. 2016; Acheson et al. 2019). All of them are outer membrane porins characterized by a 16-to 18-stranded transmembrane β-barrel architecture, forming channels through which their respective EPS are secreted to the extracellular milieu. The pore diameter varies depending on the porin but has been estimated to be around 15 Å diameter within the membrane plane in some cases, such as in BcsC (Acheson et al. 2019). In turn, TolC proteins consist of homotrimeric 12-stranded β-barrel structures (four β-strands from each protomer), forming a channel–tunnel with an average accessible interior diameter of about 19.8 Å (30 Å backbone-to-backbone) spanning the outer membrane and most of the tunnel (Koronakis et al. 2004). This suggests a degree of lumen compatibility that could potentially accommodate the extrusion of a β-glucan such as MLG. However, we also cannot rule out that TolC’s role in MLG production is indirect, e.g., through the secretion of another macromolecule necessary for the proper assembly of MLG outside the cell. In this sense, TolC is, for example, required for the secretion of ExpE1, a critical factor for efficient EPS-II synthesis or secretion in *S. meliloti* (Cosme et al. 2008). There is little doubt that future experiments capable of confirming or refuting the direct involvement of TolC in the export of certain EPS, such as MLG, could represent a significant step forward in our understanding of this broadly conserved family of bacterial proteins.

Whether directly or indirectly, our results undoubtedly demonstrate, not only that TolC is essential for MLG production in *S. meliloti*, but also that its heterologous overexpression, together with the biosynthetic genes (*bgsBA*) and the source of its allosteric activation by c-di-GMP (*pleD\**), greatly enhances MLG production. This was confirmed both with an indirect quantification by an approximately 25% increase in free glucose equivalents released after MLG lichenase digestion, and—more importantly—directly by an almost tenfold increase in the amount of MLG obtained under biotechnological fermentation conditions. The successful “bench to market” transfer of knowledge of a novel bacterial biopolymer like MLG critically depends on our capacity to scale up its production from the milligram quantities required for initial characterization to the amounts necessary for comprehensive evaluation of its biotechnological potential. This scale-up must be rationally carried out, prioritizing cost-effective and sustainable production conditions. Achieving up to 10 g/L of MLG through the specific genetic modifications described here under bioreactor conditions provides a robust foundation for future optimization of the physicochemical characteristics of the scale up fermentation process (e.g., carbon and nitrogen sources, aeration and temperature).

Additionally, the construction in this work of the plasmid pJBpleD*9091C, which carries all the known genetic factors required for MLG production, has not only enabled an increase in yield but also opens the possibility of using diverse natively non-producing bacterial chassis that may be more suitable for MLG biosynthesis. The use of alternative microbial platforms is crucial for coupling the synthesis of this biopolymer to other industrial by-products, enabling its generation from sustainable carbon and nitrogen sources that could reduce operational expenses and promote a circular economy. Future studies will evaluate this promising biotechnological approach by employing bacterial hosts beyond rhizobia. The goal is to leverage their metabolic flexibility to minimize production costs by utilizing diverse low-cost and renewable industrial by-products as feedstock.

## Conclusions

This study identifies TolC as an essential factor for MLG production in *S. meliloti*, expanding the current understanding of the molecular machinery involved in bacterial β-glucan biosynthesis. Our results support a model in which TolC participates in a tripartite complex with BgsA and BgsB, likely facilitating polymer export across the outer membrane. Importantly, overexpression of *tolC* in combination with *bgsBA* and *pleD*\* significantly enhances MLG production, enabling yields compatible with industrial-scale applications. Moreover, the successful transfer of this genetic module to non-native hosts demonstrates its versatility and opens new avenues for engineering alternative microbial platforms for MLG production. These findings provide a solid foundation for future optimization strategies aimed at sustainable and cost-effective biopolymer manufacturing.

## Author contributions

Conceptualization: DPM, JSP; Funding acquisition and Project Administration: DPM, JSP; Investigation and Methodology: LR-S, PP, JP, JLL, SM; Writing – original draft: LR-S, PP, JLL, DPM; Writing – review & editing: JSP.

## Acknowledgments

We would like to thank Miguel Redondo Nieto (UAM) for his invaluable assistance with the genomic localization of transposon sequences.

