## Supplemental Material for "TolC is required for a Mixed-Linkage β-Glucan (MLG) biosynthesis: Engineering bacteria for MLG overproduction"

### Supplementary Material

### Table S1. Bacterial strains and plasmids used in this study

| **Strains** | **Relevant characteristic** | **Reference or source** |
|---|---|---|
| ***Sinorhizobium meliloti*** | | |
| 2011 | Wild-type (*expR*$^-$) | (Meade and Signer 1977) |
| 1021 | Sm$^r$ derivative of 2011 (*expR*$^-$) | (Meade et al. 1982) |
| 8530 | *expR*$^+$ derivative of 1021 | (Glazebrook and Walker 1989) |
| 8530 Δ*bgsBA* | Derivative of 8530 containing a deletion on *Smb20390-20391* genes (*bgsBA*) | (Pérez-Mendoza et al. 2015) |
| IBR505 | Derivative of 8530 containing a deletion in *bgrV*; Sm$^r$ | (Baena et al. 2019) |
| IBR606 | Derivative of 8530 containing deletions in EPSI, EPS II and APS genes | This work |
| IBR 607 | Derivative of IBR505 containing deletions in EPSI, EPS II and APS genes | This work |
| 1021 *tolC* | Derivative of 1021, containing insertional mutation in *tolC* | (Cosme et al. 2008) |
| 2011 *tolC* | Derivative of 2011, containing insertional mutation in *tolC* | (Cosme et al. 2008) |
| ***Rhizobium etli*** | | |
| CFN42 | Wild-type; Nx$^r$ | (Quinto et al. 1985) |
| Δ*celAB* | Derivative of CFN42 containing a deletion on *RHE_CH01542*-CH01543 (*celAB*) | (Pérez-Mendoza et al. 2022) |
| Δ*celAB* Δ*bgsA* | Derivative of Ret Δ*celAB*, containing a deletion in RHE_PE00363 (*bgsA*); Nx$^r$ | (Pérez-Mendoza et al. 2022) |
| ***Escherichia coli*** | | |
| DH5α | *supE44, ΔlacU169, Φ80, lacZΔM1, recA1,endA1, gyrA96, thi1, relA1, 5hsdR171* | (Hanahan 1983) |
| NovaBlue | *endA1 hsdR17* (*rK12– mK12+*) *supE44 thi-1 recA1 gyrA96 relA1 lac F'*[*proA+B+ lacIqZΔM15::Tn10*];Tc$^r$ | Novagen |

| | | |
|---|---|---|
| β2163 | (F−) RP4-2-Tc::Mu Δ *dapA*::(*erm-pir*);$Km^r$, $Em^r$, $DAPA^{aux}$ | (Demarre et al. 2005) |
| S17.1 | thi, pro, recA, hsdR, hsdM, Rp4Tc::Mu, Km::Tn7; $Sm^R$, $Sp^R$, $Tmp^R$ | (Simon et al. 1983) |
| ***Plasmids*** | | |
| pJB3Tc19 | Cloning vector, $P_{lac}$ promoter; $Ap^r$, $Tc^r$ | (Blatny et al. 1997) |
| pJBpleD* | pJB3Tc19 derivative harbouring a 1423 bp XbaI/EcoRI fragment containing *pleD** from *Caulobacter crescentus*; $Ap^r$, $Tc^r$ | (Pérez-Mendoza et al. 2014) |
| pJBpleD*9091 | pJB3Tc19 derivative bearing a EcoRI fragment containing SMb20390 (*bgsB*) and SMb20391 (*bgsA*) and 1423 pb XbaI/EcoRI fragment containing *pleD**; $Ap^r$, $Tc^r$ | (Pérez-Mendoza et al. 2015) |
| pJBpleD*9091C | pJBpleD9 derivative bearing a 4976 bp XbaI fragment containing *bgsBA* and tolC; $Ap^r$, $Tc^r$ | This work |
| pJBpleD*lacI$^q$ | pJBpleD* derivative containing a *lacI$^q$* allele | This work |
| pSUP2021 | pSUP202 derivative harbouring the Tn5 transposon, $Cm^r$, $Ap^r$, $Tc^r$ | (Simon et al. 1983) |
| pSAM_Rl | pSAM_Km derivative with *B. thetatiotamicron rpoD* promoter replaced with *R. leguminosarum* 3841 *rpoD* promoter region, $Ap^r$, $Km^r$ | (Perry and Yost 2014) |
| pQE-80L | Medium-copy expression vector, IPTG-inducible T5 promoter, $Ap^r$ for N-terminus His6-tag fusions; $Ap^R$ | Qiagen® |

### Table S2. Primers used in this work

| Primers | Product | Sequence (5´-3´) | Used in |
|---|---|---|---|
| TolC-F<br>TolC-R | 1457 bp | GGATCCCCGCCAAACGGTCTGC<br>GGATCCTGTCGAACTGGCTGAGGG | Amplification of *tolC* from *Ensifer meliloti* |
| EcoRItolC-F<br>HindIIItolC-R | 2680 bp | TTAAAGAATTCAAAGCTCCTCCAGCTCGC<br>ATTTAAAGCTTCGGCTCCTGCTTGATCCC | Amplification of *tolC* and 709 bp and 564 bp upstream and downstream, respectively |
| Tn5F1<br>Tn5F2 | - | TGCATGGCTTCACTACCC<br>GAGAACACAGATTTAGCCCA | Forward primers for random PCR on Tn5-harbouring genomic DNA (Tn5F1 for round 1 and Tn5F2 for round 2) |

| | | | |
|---|---|---|---|
| SAMIRLSEC<br><br>IRL3 | - | CGAAACGATCCTCATCCT<br><br>TTCGCTTGCTGTCCATAAAACCGCCC | Forward primers for random PCR of IRL on SAM-harbouring genomic DNA (SAMIRLSEC for round 1 and IRL3 for round 2) |
| IRR2<br><br>SAMIRRSEC | - | GGACGCCCGCCATAAACTGCC<br><br>CAGAAGGCCATCCTGAC | Forward primers for random PCR of IRR on SAM-harbouring genomic DNA (IRR2 for round 1 and SAMIRRSEC for round 2) |
| JAMS_1R<br><br>JAMS_2R | - | GGCCACGCGTCGACTAGTACNNNNNNNNNNACGCC<br><br>GGCCACGCGTCGACTAGTAC | Reverse primers for random PCR of IRR/IRL on Tn5/SAM-harbouring genomic DNA (JAMS_1R for round 1 and JAMS_2R for round 2) |
